# V2b neurons act via multiple targets in spinal motor networks

**DOI:** 10.1101/2022.08.01.502410

**Authors:** Mohini Sengupta, Martha W. Bagnall

**Affiliations:** Washington University School of Medicine, Department of Neuroscience, St Louis, MO, USA

## Abstract

Within the spinal cord, interneurons shape motor neuron activity. These interneurons can project over long distances in the longitudinal axis, but systematic mapping of their connectivity has been limited. In this study, using larval zebrafish, we mapped local and long-range connectivity of a cardinal spinal population, the Gata3^+^ V2b class. V2b neurons are inhibitory and project ipsilateral, descending axons. We show that V2b neurons are active during fictive swimming, slightly leading the motor burst. Via optogenetic mapping of output in the rostrocaudal axis, we demonstrate that V2b neurons robustly inhibit motor neurons and other major spinal interneuron classes, including V2a, V1, commissural neurons and other V2b neurons. V2b inhibition is patterned along the rostrocaudal axis, providing long-range inhibition to motor and V2a neurons but more localized innervation of the V1 class. Furthermore, these results provide the first demonstration of reciprocal V1/V2b inhibition in axial circuits, potentially representing an ancestral motif of the limb control network.

## Introduction

In vertebrates, locomotor movements are executed by intrinsic networks in the spinal cord^1–3^. Spinal networks comprise excitatory and inhibitory interneurons that connect with each other and with motor neurons to orchestrate different movements^3,4^. Across vertebrates, neuronal genetic identity, morphology, and neurotransmitter expression remain remarkably conserved, making it convenient to study and apply knowledge across species^3,5^. Behaviorally, coordination along the rostrocaudal axis during locomotion is also a conserved feature^6–9^, yet mechanisms that facilitate this coordination are not understood. A primary reason for this gap is the lack of knowledge on long range connectivity of neurons. In our previous work, we revealed that spinal V1 neurons inhibited different post-synaptic targets locally and distally, revealing significant variations in longitudinal connectivity and function^10^. It remains unclear whether these variations in rostrocaudal connectivity are specific to V1 neurons or a more general property of spinal circuits.

Spinal V2b neurons are a major inhibitory population marked by expression of the Gata3 transcription factor across vertebrates^11–13^. In mice^14^ and zebrafish^13^, V2b neurons project ipsilateral descending axons spanning several segments, but apart from contacting motor neurons and other V2b neurons^13^, postsynaptic partners of this population remain unknown. In mice, V2b neurons have been identified as a subset of long descending propriospinal neurons that connect cervical and lumbar segments^14^ and are thought to implement inter-limb coordination during locomotion^15^.

Functionally, V2b neurons have primarily been implicated in limb movements, where together with V1 neurons, they enforce flexor-extensor alternation^16,17^. However, the V2b population is present not only in spinal segments that control limb motor neurons but all along the spinal cord in both mice^18^ and zebrafish^13^, suggesting an ancestral role in locomotor coordination. In these axial circuits, some evidence suggests that V2b neurons can serve as locomotor brakes^13^. Bilateral optogenetic activation of V2b neurons in larval zebrafish reduced frequency of tail movements, whereas suppression of V2b neurons led to accelerated swimming.

In this study, we analyzed the recruitment pattern and longitudinal connectivity of V2b neurons in axial motor networks of larval zebrafish. We show that V2b neurons are active during locomotion, with a modest phase lead in the swim cycle. Via circuit mapping, we demonstrate that V2b neurons inhibit motor neurons as well as other cardinal excitatory and inhibitory spinal populations, including V1 neurons. Moreover, V2b inhibition is structured in the rostrocaudal axis, providing a graded input to motor and V2a neurons but more localized input to V1 neurons. Combined, these results indicate that V2b neurons have a multifaceted role in axial motor control, and their reciprocal connectivity with V1 neurons forms a motif that may recur in limb control circuits.

## Materials and Methods

### Animals

Adult zebrafish (Danio rerio) were maintained at 28.5°C with a 14:10 light:dark cycle in the Washington University Zebrafish Facility following standard care procedures. Larval zebrafish, 4–6 days post fertilization (dpf), were used for experiments and kept in petri dishes in system water or housed with system water flow. Animals older than 5 dpf were fed rotifers or Gemma dry food daily. All procedures described in this work adhere to NIH guidelines and received approval by the Washington University Institutional Animal Care and Use Committee.

### Transgenic Fish Lines

For recruitment studies in Figure 1, V2b neurons were targeted in the stable *Tg (Gata3:LoxP-dsRed-LoxP:GFP)^nns53Tg^* (ZDB-ALT-190724-4)^13^. For all connectivity experiments, the stable transgenic line *Tg(Gata3:Gal4;UAS:CatCh)^stl602^* (DB-ALT-201209-12)^13^ generated by Tol2 mediated transgenesis previously in our lab was used. For targeting V2a and V1 neurons, the *Tg(vsx2:loxP-DsRed-loxP-GFP) ^nns3Tg^* (ZDB-ALT-061204-4)^19^ and *Tg(En1: LoxP-dsRed-LoxP:DTA) ^[nns55Tg](https://zfin.org/ZDB-ALT-191030-2)^* (ZDB-ALT-191030-2)^20^ lines, respectively, were crossed to *Tg(Gata3:Gal4;UAS:CatCh)* to generate double transgenics. Secondary motor neurons were targeted in part using the *Tg(mnx:pTagRFP) ^stl603^* line created in the lab.

**Figure 1:**
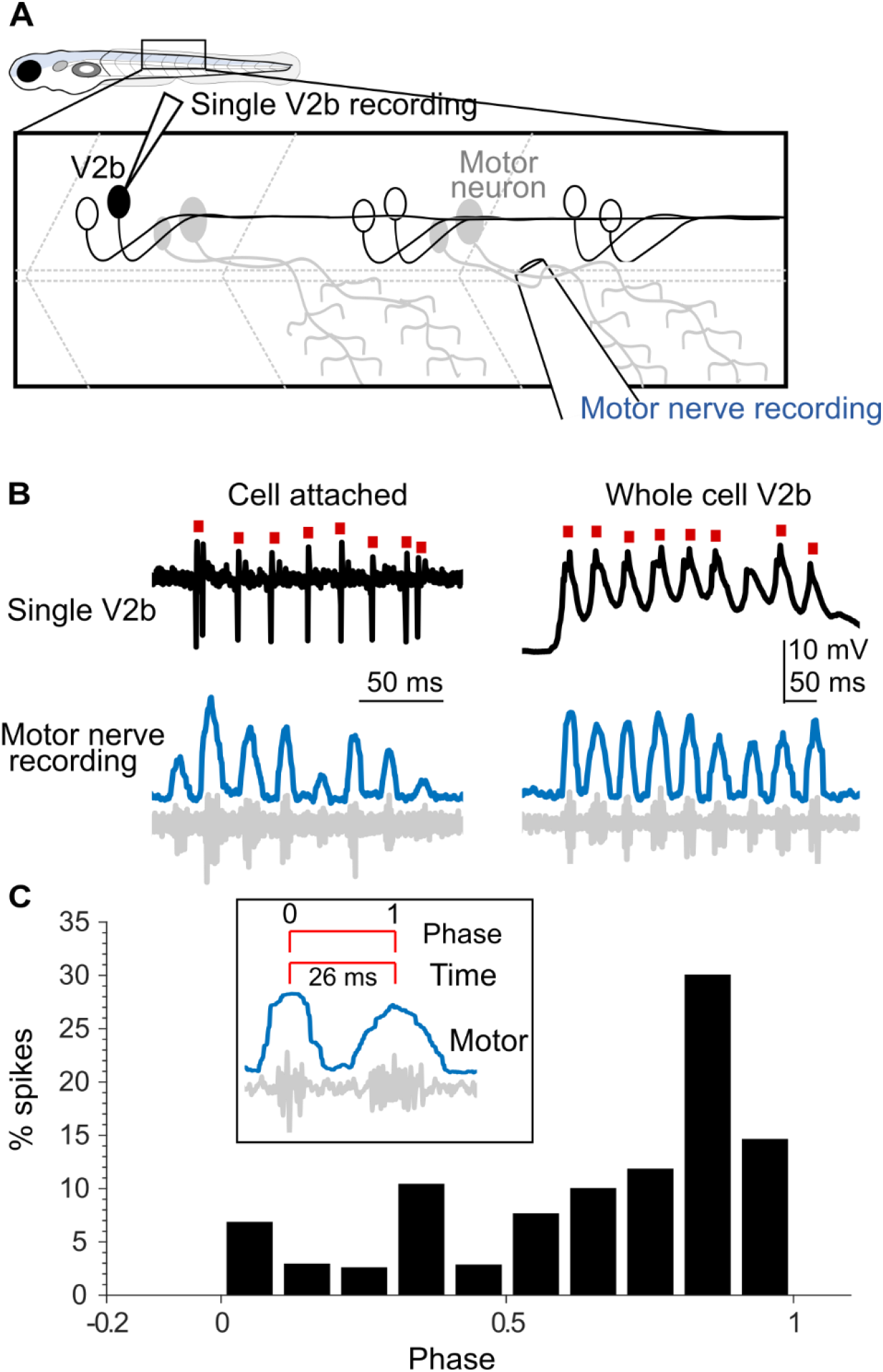
Activity of V2b neurons during fictive locomotion. A. Schematic of the experimental set-up showing simultaneous intracellular recording from single V2b neurons and extracellular recording of the motor nerve. B. Representative traces showing V2b spikes (top, black) recorded in cell attached (left) or whole cell (right) mode during motor activity (bottom, grey). Red squares mark V2b spikes. The standard deviation over a sliding 10 ms window was used to determine the midpoint of motor bursts (blue trace). C. Histogram of V2b spike timing with respect to normalized phase of motor activity. Inset: Illustration for timing and phase relation between successive motor bouts. N= 685 swim cycles from 11 neurons.

### Electrophysiology

4–6 dpf larvae were immobilized with 0.1% α-bungarotoxin and fixed to a Sylgard lined petri dish with custom-sharpened tungsten pins. One muscle segment overlaying the spinal cord was removed at the mid-body level (segments 9–13). The larva was then transferred to a microscope (Nikon Eclipse E600FN) equipped with epifluorescence and immersion objectives (60X, 1.0 NA). The bath solution consisted of (in mM): 134 NaCl, 2.9 KCl, 1.2 MgCl_2_, 10 HEPES, 10 glucose, 2.1 CaCl_2_. Osmolarity was adjusted to ~295 mOsm and pH to 7.5. For recording V2b spiking during swims, a combination of cell attached and whole cell patch clamp recordings were obtained from V2b neurons. Patch pipettes (7–15 MΩ) were either filled with extracellular saline (cell attached) or patch internal (whole cell) solution and targeted to a V2b neuron. After formation of a gigaohm seal, extracellular spikes were recorded in cell attached mode. To obtain whole cell recordings, brief suction pulses were used to break into the cell. Spiking was recorded in current clamp mode using the following patch internal solution: (in mM) 125 K gluconate, 2 MgCl_2_, 4 KCl, 10 HEPES, 10 EGTA, and 4 Na_2_ATP. Ventral root recordings were obtained 2-3 segments caudal to the recorded V2b, using suction electrodes (diameters 20-50 μm). Fictive swimming (~20-50 Hz) was elicited by either a brief electric stimulus to the tip of the tail (10-20 V for 0.2-1 ms)^20^ or a visual stimulus of blue and white moving gratings (spatial width of 1 cm), projected 1 cm away, and moved at 1 cm/s^21^.

For mapping connectivity to target neurons, larvae were immobilized and dissected as before. Whole cell patch clamp recordings were made from neurons in voltage clamp mode using the following patch internal solution: (in mM) 122 cesium methanesulfonate, 1 tetraethylammonium-Cl, 3 MgCl_2_, 1 QX-314 Cl, 10 HEPES, 10 EGTA, and 4 Na_2_ATP. APV (10 μM) and NBQX (10 μM) were added to the bath to block glutamatergic transmission. For all patch internal solutions, pH was adjusted to 7.5 and osmolarity to 290 mOsm. Additionally, for identifying primary motor neurons, commissural neurons and sensory populations, sulforhodamine 0.02% was included in the patch internal to visualize morphology of recorded cells post-hoc. Recordings were acquired using a Multiclamp 700B amplifier and Digidata 1550 (Molecular Devices). Signals were filtered at 2 kHz and digitized at 100 or 50 kHz. For IPSCs, cells were recorded at +0.3 mV (after correction for liquid junction potential of 14.7 mV).

### Optogenetic Stimulation

A Polygon 400 or 1000 Digital Micromirror Device (Mightex) was used to deliver optical stimulation. The projected optical pattern consisted of a 4×4 grid of 16 squares. Each square in the grid approximately measured 20 μm × 20 μm and covered 0-6 cells depending on position. One full stimulus pattern consisted of an ordered sequence of each of the 16 squares sequentially. The 16^th^ square in most cases spilled out to neighboring segments or out of the spinal cord and hence was not included in the data. For each small square, illumination consisted of a 20 ms light pulse (470 nm) at 50% intensity (4.6-5.2 μW under 60X, 1.0 NA). The sequence was triggered using a TTL pulse from the Digidata to synchronize the stimulation with electrophysiology. The objective was carefully positioned over a single spinal segment prior to stimulus delivery; for each new segment, the stage was manually translated and repositioned. V2b spiking reliability was measured by delivering multiple trials to a selected square that had evoked spiking on the first trial. For the high frequency stimulation in Fig. 3, a single square was illuminated with a 20 Hz train of five 20 ms pulses.

### Analysis

Electrophysiology data were imported into Igor Pro 6.37 (Wavemetrics) using NeuroMatic^22^. Spikes and IPSCs were analyzed using custom code in Igor and MATLAB. For analyzing motor recordings from the ventral nerve (Fig 1), the raw swim signal (Fig. 1B, grey traces) was converted to the standard deviation signal over a sliding window of 10 ms (Fig. 1B, blue traces). Peaks from this SD signal, corresponding to the midpoint of motor bouts, were calculated using custom written MATLAB codes. V2b spike times were normalized relative to the interbout duration. Histograms of these spike times were normalized for each cell and then pooled together (from 12 cells) for the summary plot shown in Fig. 1C. For connectivity experiments (Fig 3-7), data were analyzed as previously reported^10^. Briefly, charge transfer for the evoked response was calculated by integrating the current in a 50 ms window from the onset of the optical stimulus (Evoked) and subtracting this from Control 1, a similar integral over a 50 ms window before the optical stimulus. This was done to account for spontaneous activity. To calculate background noise values, a similar integral for a different 50 ms window at the end of the recording (Control 2) was subtracted from Control 1. Both the charge transfer of the evoked response and background noise were summed across the 16 squares for each segment.

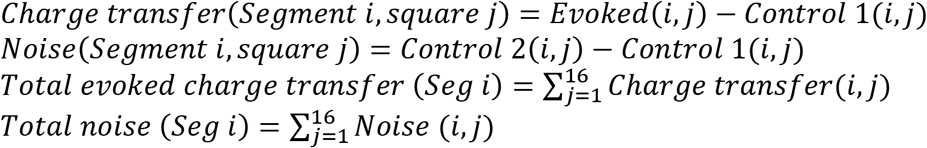

For statistical comparisons, *Total evoked charge transfer (Seg i)* was compared to *Total noise (Seg i)* for each target population using the Wilcoxon Sign Rank Test (p<0.05). Statistical tests were performed using MATLAB (R2020b, MathWorks). Due to the non-normal distribution of physiological results, including spiking and IPSC charge transfer, we used nonparametric statistics and tests.

Peak amplitudes of IPSCs were calculated as the maximum value of the charge transfer trace. Conductances were calculated as peak amplitude / driving force (75 mV). Input resistance was measured by an average of small hyperpolarizing pulses.

## Results

### V2b activity during fictive swimming

Spinal interneurons in the ventral horn that participate in generating rhythm or pattern of motor activity show stereotypic firing patterns during movements^20,23,24^. To determine the firing patterns of V2b neurons, we recorded from identified neurons in the transgenic *Tg(Gata3:lox-dsred-lox:GFP)* line while simultaneously monitoring fictive motor activity from ventral nerve responses in paralyzed zebrafish larvae at 4-6 dpf (Fig. 1A). Fictive locomotion (20-50 Hz) was evoked either by a brief electric shock to the tail or a visual stimulus of moving bars. Over half of V2b neurons (20 out of 32 neurons, 62.5%) did not exhibit any spiking during fictive swims. Among V2b neurons that did fire during motor episodes, spiking rates were similar between cell attached (Fig. 1B, left) and whole cell (Fig. 1B, right) recordings and hence these were pooled for subsequent analysis. To investigate the phase of V2b spiking relative to the swim cycle, we constructed histograms of V2b spike times normalized with respect to motor activity (Supplementary Fig. 1). Most V2b neurons (7 out of 11), exhibited spiking that was coupled to a specific phase of motor activity (Rayleigh’s Test). The remaining V2b neurons (4 out of 11) exhibited fewer spikes that were more distributed, lacking a clear phase relationship to motor activity. The population data is summarized in Fig. 1C. Taken together, the highest frequency was centered at 0.85, indicating that most V2b activity coincides with the rising phase of on cycle excitation and motor neuron spiking. This is also evident from representative traces in Fig. 1B. To determine how V2b-mediated inhibition may influence the spinal circuit, we next engaged in systematic mapping of the postsynaptic partners of V2b neurons.

### Optogenetic assisted mapping of V2b connectivity

To map V2b connectivity along the rostrocaudal axis, we utilized the high throughput technique of optogenetic assisted circuit mapping^25^. We used the transgenic line *Tg(Gata3:Gal4;UAS:CatCh)*, in which V2b neurons expressed a calcium permeable variant of Channelrhodopsin, CatCh^26^ (Fig. 2A, schematic). We first calibrated the efficacy and specificity of the optical stimulus for activating V2b neurons restricted to a single segment. A 4×4 square grid pattern was projected approximately over a single spinal segment (Fig. 2B). Each square in this grid measured 20 × 20 μm and was illuminated in a sequential order with a 20 ms light pulse. V2b neurons recorded in current clamp mode exhibited a mean resting membrane potential of −74.3 ± 2.37 mV. Direct illumination on the soma (black dot) or nearby elicited robust spiking in V2b neurons (Fig. 2C, red traces). V2b neurons project descending axons in the R-C axis, extending an average of seven segments^13^. We tested whether the optical stimulus could elicit antidromic spikes when the axon was illuminated by translating the stimulus to neighboring rostral and caudal segments while recording V2b somatic responses (Fig. 2D). Optical stimulation outside of Segment 0 (segment in which V2b is being recorded) did not evoke any appreciable activity (Fig. 2D, E, N=10 neurons). V2b neurons also showed high reliability of spiking to repeated presentations of the optical stimulus, with a mean reliability of 75% ± 12.1 (Fig. 2F, N=10 neurons). Together, these data show that the grid optical stimulus activated V2b neurons in a spatially restricted manner and therefore can be utilized for mapping V2b connectivity to different targets along the R-C axis.

**Figure 2:**
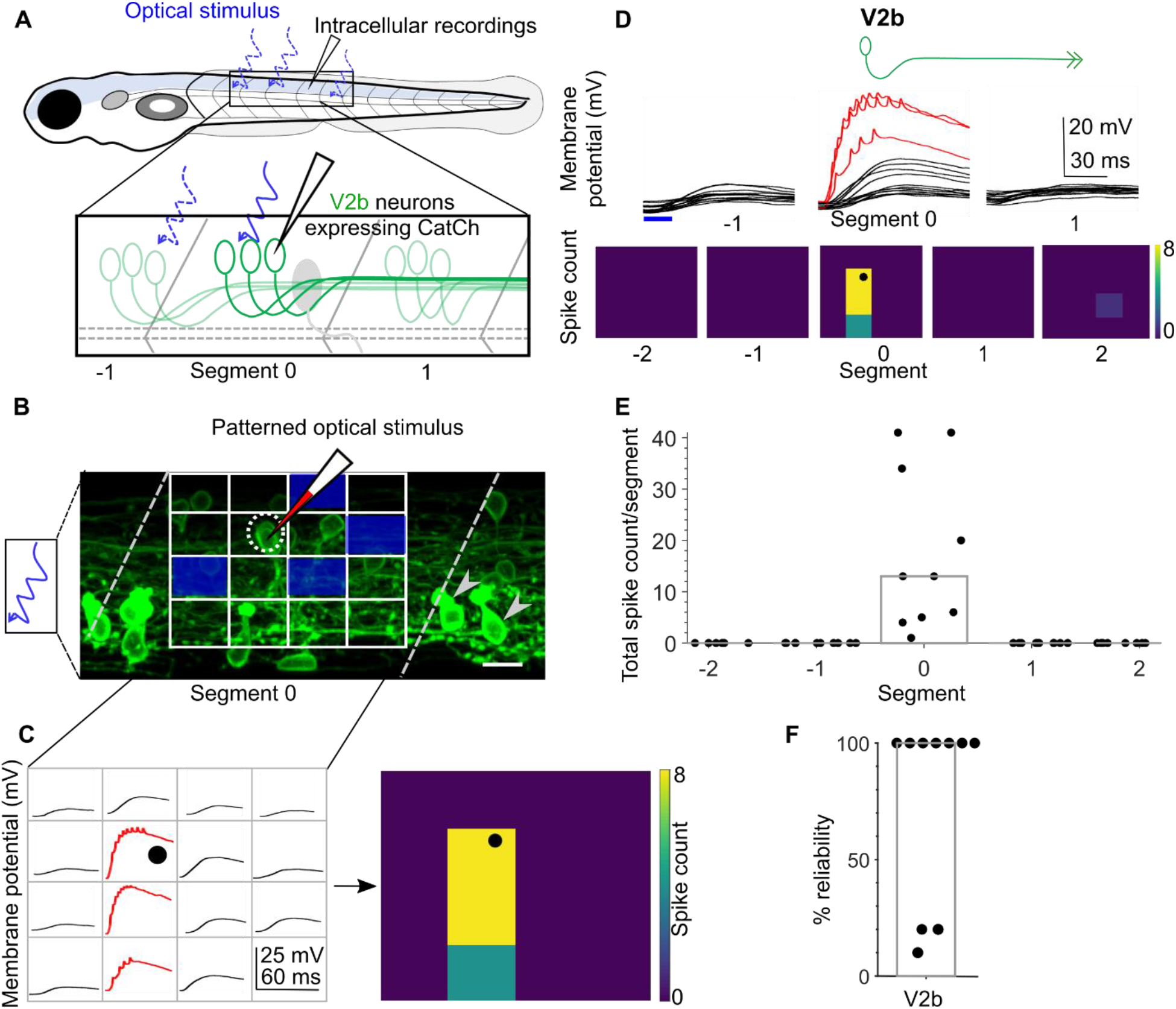
Calibration of V2b activation by patterned optical stimuli. A. Schematic of the experimental set-up showing targeted intracellular recording and optical stimulation in *Tg(Gata3:Gal4,UAS:CatCh)* animals. B. Schematic of the patterned optical stimulus. A 4×4 grid was overlaid on approximately one spinal segment and each small square in the grid (blue square) was optically stimulated in sequence with a 20 ms light pulse. Position of the recorded cell is outlined in a grey dotted circle. Arrowheads show CSF-cNs also labeled in this line. C. Intracellular recordings elicited from optical stimulation in each grid square (left). Spiking is denoted in red. Same data shown as a heat map superimposed on the stimulus grid (right). Position of the recorded cell is indicated with a black dot. D. V2b responses evoked by optical stimuli in segments rostral or caudal to the recorded neuron. Representative traces of activity (top) and spike count (bottom) of the same V2b neuron while the optical stimulation was translated along the rostro-caudal axis. Red traces denote spiking. E. Quantification of spiking in V2b neurons as the optical stimulus is presented along the rostro-caudal axis. N = 10 neurons. Bar indicates median value. F. Reliability of spiking in these neurons with multiple trials of the same optical stimulus. N=10 neurons. Bar indicates median value.

### V2b neurons robustly inhibit motor neurons up to long distances

In mice, V2b neurons in lumbar segments make anatomical contacts to motor neurons, preferentially those controlling extensors^16^. In larval zebrafish, physiological studies show that V2b neurons inhibit both fast, primary motor neurons and slow, secondary motor neurons^13^, but it is not known if this connectivity is only local or extends over many segments. To investigate this, we recorded intracellularly from primary motor neurons in the *Tg(Gata3:Gal4;UAS:CatCh)* line (Fig. 3A). Primary motor neurons were identified in bright field with their characteristic large, laterally positioned soma and post hoc dye fill showing extensive muscle innervation^27^. Cells were recorded using a cesium internal, held at 0 mV and bathed in glutamate receptor blockers (NBQX, APV) to isolate inhibitory post synaptic currents (IPSCs). The grid optical stimulus was delivered at a single segment each time and translated rostrally for 7 segments to cover the full descending axon length of V2b neurons.

**Figure 3:**
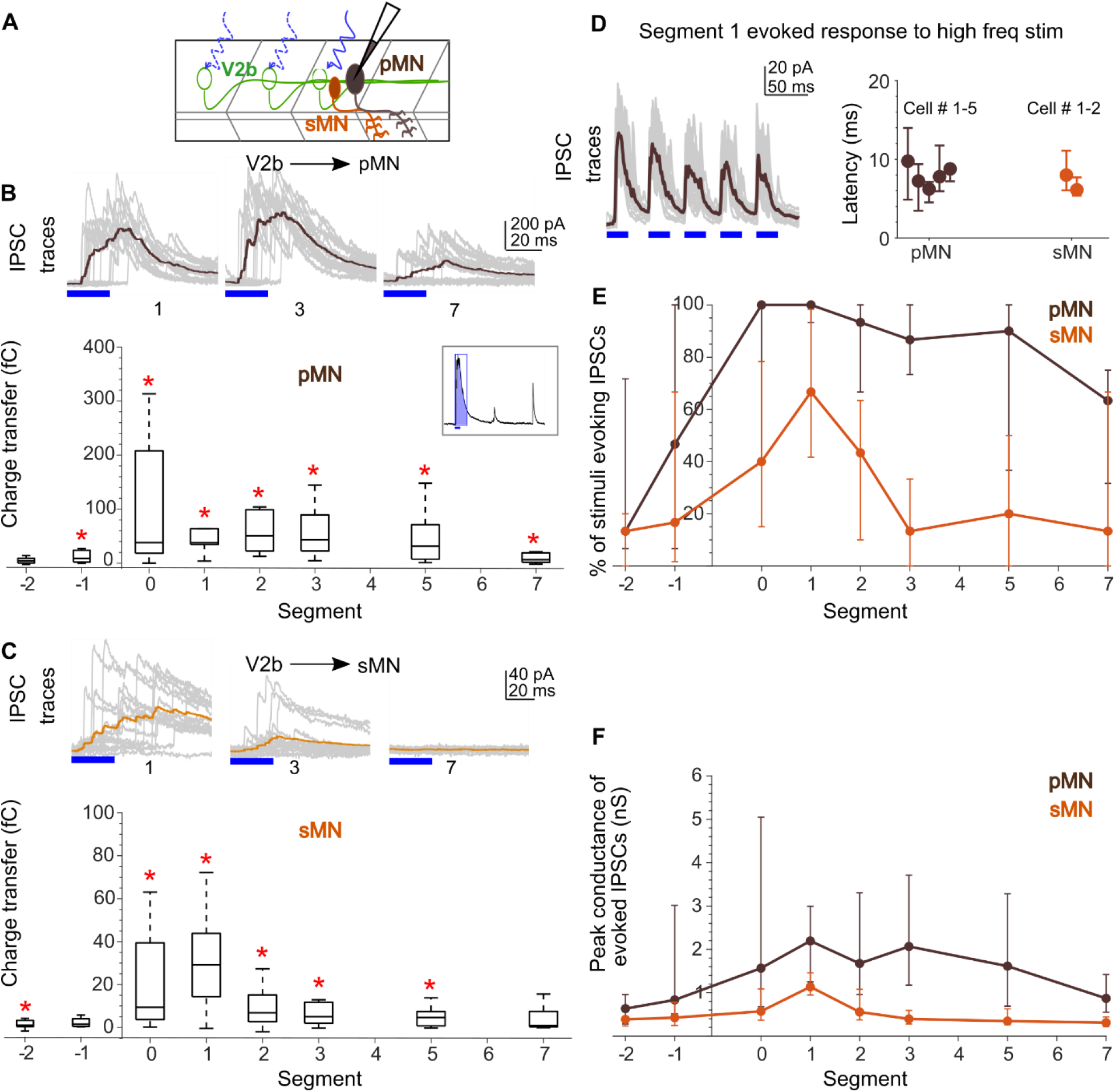
Motor neurons receive short- and long-range inhibition from V2b neurons. A. Schematic of the experimental design showing intracellular recordings from primary (brown) and secondary (orange) motor neurons paired with optical stimulation of CatCh+ V2b neurons (green) along the rostrocaudal axis. B. Top: Representative overlay of 15 traces of IPSCs recorded in a primary motor neuron (pMN) during illumination of segments 1, 3, and 7 rostral to the recorded neuron soma. Colored trace represents mean. Optical stimulus is represented as a blue bar. Bottom: Box plots showing total charge transfer per segment (inset, illustration) recorded in primary motor neurons. In this and subsequent Figures, the box shows the median, 25^th^, and 75^th^ percentile values; whiskers show +/–2.7σ. Red asterisks mark distributions that were significantly different from noise (*p* < 0.01, Wilcoxon Sign Rank Test). C. Same as in B for secondary motor neurons (sMNs). D. Left: Representative overlay of 10 traces of IPSCs recorded in primary motor neurons (pMNs) during a 20 Hz optical stimulus train (5 pulses, 20 ms) on segment 1. D. Right: Median latency of the first IPSC for each of the five pulses in the train stimulus for each neuron. Error bars represent 25^th^ and 75^th^ percentiles. Dots represent Cell # 1-5 for pMNs (brown) and 1-2 for sMNs (orange). E, F. Comparison of the percent of squares in the optical stimulus grid that evoked IPSCs (E) and the peak conductance of IPSCs (F) in primary (brown) and secondary (orange) motor neurons. Here and in subsequent Figures, circles represent median values and error bars indicate the 25^th^ and 75^th^ percentiles. N= 10-25 neurons for pMNs and 7-11 neurons for sMNs.

Primary motor neurons recorded in this configuration exhibited IPSCs when V2b neurons were optically activated nearby or at long range (Fig. 3B). To quantify these inputs, we calculated the charge transfer of evoked IPSCs (Fig. 3B, inset) for each segment relative to noise (see Methods). Primary motor neurons received significant V2b-mediated inhibition even up to seven segments distant from the site of stimulation (N=10-25 neurons; Wilcoxon Sign Rank Test, *p* < 0.01). We next performed similar mapping from V2b neurons to secondary motor neurons, which were identified either genetically in the double transgenic *Tg(Gata3:Gal4;UAS:CatCh ; mnx:ptag:RFP)* line or by post hoc dye label. Secondary motor neurons also received local and long-range inhibition from V2b neurons (Fig. 3C). Charge transfer at Segments 1 through 5 was significantly above noise levels (Fig. 3C, lower panel, N=7-11 neurons; Wilcoxon Sign Rank Test, *p* < 0.01) but decreased to insignificant at 7 segments away.

V2b neurons in zebrafish have purely descending axons, and therefore are not expected to inhibit neurons in the rostral direction. Consistent with this expectation, we saw little to no evoked IPSCs during delivery of optical stimuli caudal to the recorded segment (Segments −1, −2) (Fig. 3B, C). Because our transgenic line also labels CSF-cNs (Fig. 2B, grey arrowheads), which have purely ascending axons^28,29^ and are known to selectively inhibit one of the four primary motor neurons^30^, this low-frequency connectivity likely results from activation of CSF-cNs.

To verify that evoked IPSCs were monosynaptically generated, we delivered a train of optical stimuli (five 20 ms pulses at 20 Hz). IPSCs followed the train with consistent latency and low jitter, consistent with monosynaptic connectivity (Fig. 3D).

Finally, we asked if V2b connectivity to motor neurons showed differences in number and/or strength of connections with distance. As a proxy for number of connections, we quantified the fraction of small squares in the 4×4 grid stimulus in each segment that successfully evoked IPSCs. To quantify IPSC amplitude, we calculated the average maximum conductance for each segment. Interestingly, connectivity rates remained high along the longitudinal axis, with 63% of squares evoking IPSCs onto primary motor neurons even at 7 segments away (Fig. 3E, N=10-25 pMNs). However, the amplitude of these V2b contacts onto primary motor neurons tapered off at long ranges (Fig. 3F). In contrast, for secondary motor neurons, both the number of V2b contacts (80% reduction) as well as the strength of connections (73% reduction) declined from Segment 1 to Segment 7 (Fig. 3E and F, N=7-11 sMNs). Taken together, these data show that V2b neurons monosynaptically inhibit both primary and secondary motor neurons locally and at long distances. The strength of V2b-mediated inhibition progressively diminishes with distance but extends further for primary than for secondary motor neurons.

### V2b neurons inhibit Chx10^+^ V2a neurons over long ranges

Spinal V2a neurons are a major source of excitatory drive to motor neurons^31–35^. These neurons participate in rhythm generation and are indispensable for motor functions^34,36^. Therefore, we next examined V2b-mediated inhibition of the V2a population along the longitudinal axis. V2a neurons were targeted using the double transgenic *Tg(Chx10:lox-dsred-lox:GFP)*;*(Gata3:Gal4;UAS:CatCh)* line (Fig. 4A). Stimulation of V2b neurons both locally and long-range elicited inhibitory synaptic inputs in V2a neurons (Fig. 4B). Charge transfer of inhibitory currents was significantly higher than noise up to seven segments from the recording site (Fig. 4C, N=7-10 neurons; Wilcoxon Sign Rank Test, *p* < 0.01). V2a neurons did not receive any appreciable evoked synaptic input when caudal segments were illuminated (Fig. 4C, Segment −1, −2). As with primary motor neurons, the inferred number of connections was maintained along the rostrocaudal axis (26% decrease from Segment 1 to Segment 7, Fig. 4D, N=7-10 neurons) but the strength of V2b inhibitory inputs diminished gradually at long range (56% reduction from Segment 1 to Segment 7, Figure 4E). Overall, these data identify V2a population as a major novel target of V2b neurons. Furthermore, we show that V2b neurons inhibit the V2a population both locally and long-range, with a gradual reduction in strength at long distances.

**Figure 4:**
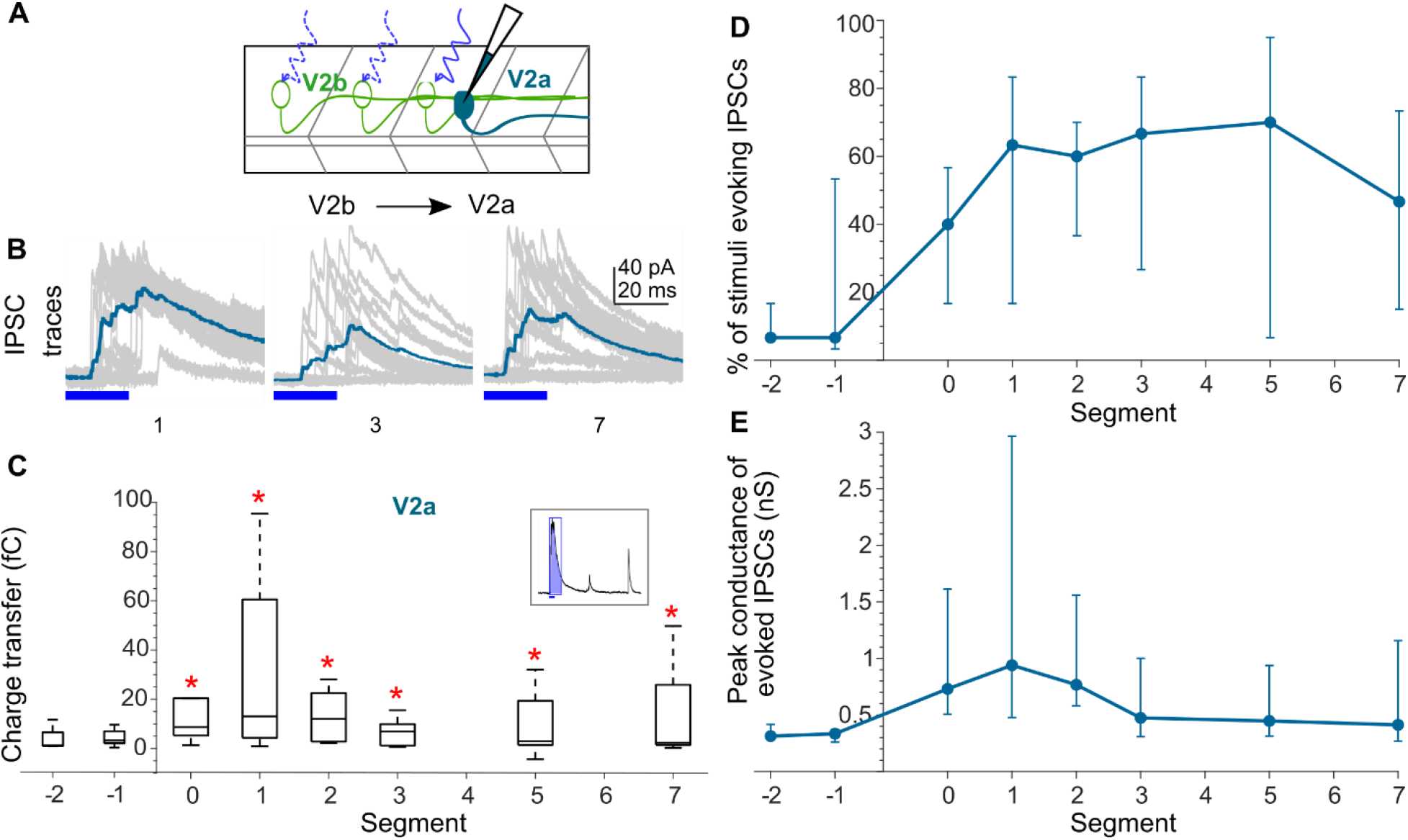
V2a neurons receive local and long-range inhibition from V2b neurons. A. Schematic of the experimental design showing intracellular recordings from V2a neurons (cyan) paired with optical stimulation of CatCh+ V2b neurons (green) along the rostro-caudal axis. B. Overlay of 15 traces recorded in a V2a neuron during illumination of segments 1, 3, and 7 rostral to the recorded neuron soma. Colored trace represents mean. C. Box plot showing total charge transfer per segment (inset, illustration) recorded in V2a neurons. D, E. Comparison of the percent of squares in the optical stimulus grid that evoked IPSCs (D) and the peak conductance of IPSCs (E) in V2a neurons. N=7-10 neurons.

### V2b neurons inhibit spinal V1 neurons locally

V1 neurons are a major inhibitory population in the spinal cord^37–39^ and share a special relationship with V2b neurons for limb control^16^. In lumbar networks that control the hind limbs, V1/V2b neurons reciprocally inhibit antagonistic motor pools, driving alternation of flexors and extensors. Flexor-extensor related Ia inhibitory interneurons have been shown to directly inhibit each other in several species^24,40,41^, including humans^42^. While Ia inhibitory neurons belong to the V1/V2b populations^43^, direct evidence of reciprocal inhibition between these two genetically defined populations has not been found. To determine if V2b neurons inhibit the V1 population, we recorded from V1 neurons identified in the *Tg(eng1b:lox-dsred-lox:DTA)* line (Fig. 5A). Activation of V2b neurons evoked significant IPSCs in V1 neurons locally, at Segments 0-3, but not long-range at Segments 5-7 (Fig. 5C, N= 8-9 neurons). Both the fraction of squares evoking IPSCs (Fig 5D) and the amplitude of evoked IPSCs (Fig 5E) were maximal at 2-3 segments from the recording site and fell off in either direction. This pattern was in contrast to the long-range V2b inhibition onto motor neuron and V2a targets. Therefore, the structure of V2b-mediated inhibition varies along the longitudinal axis to distinct downstream targets. These data are the first demonstration that V2b neurons inhibit the V1 population.

**Figure 5:**
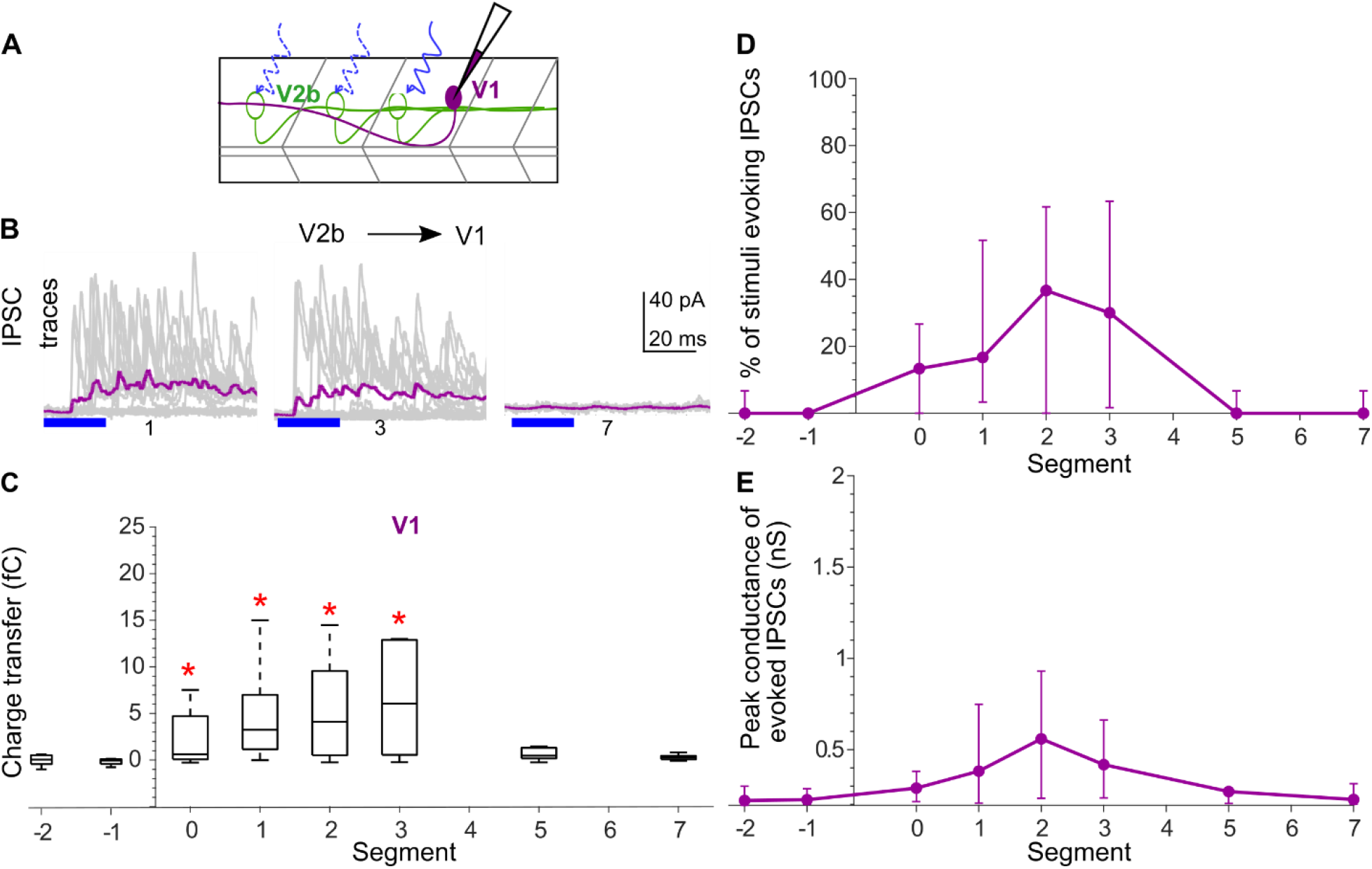
V1 neurons receive predominantly local V2b inhibition. A. Schematic of the experimental design showing intracellular recordings from V1 neurons (magenta) paired with optical stimulation of CatCh+ V2b neurons along the rostro-caudal axis. B. Representative overlay of 15 traces recorded in a V2a neuron during illumination of segments 1, 3, and 7 rostral to the recorded neuron soma. Colored trace represents mean. Duration of the optical stimulus is shown as a blue bar. C. Box plot showing total charge transfer per segment (inset, illustration) recorded in V2a neurons. D, E. Comparison of the percent of squares in the optical stimulus grid that evoked IPSCs (D) and the peak conductance of IPSCs (E) in V1 neurons. N=8-9 neurons.

### V2b connectivity to other ventral spinal populations

We next extended this map to include other spinal populations in the ventral horn. Commissural neurons in the spinal cord are a major functionally relevant group that helps secure left-right alternation^44–46^, rostro-caudal coordination^15,23^, and modulation of motor strength^47,48^. Commissural neurons comprise more than one genetically defined class (dI6, V0, and V3 neurons^45,49–52^), and also exhibit morphological variability within each class. To first determine whether V2b neurons target any commissural neurons, we performed blind recordings of neurons and classified them post hoc based on their morphology from dye fills as commissural, bifurcating and ascending neurons, likely belonging to the V0/dI6 classes. We collectively refer to these as Commissural Premotor neurons or CoPrs. CoPr neurons were located more dorsally than V3 neurons^47,48^ which therefore are unlikely to be included in this population. CoPr neurons received modest and variable local inputs from V2b neurons (Fig. 6A, N= 5-7 neurons) that were significant only at Segment 0 (Wilcoxon Sign Rank Test, *p* < 0.01). These results suggest that V2b neurons predominantly target ipsilateral pathways at long-ranges.

**Figure 6:**
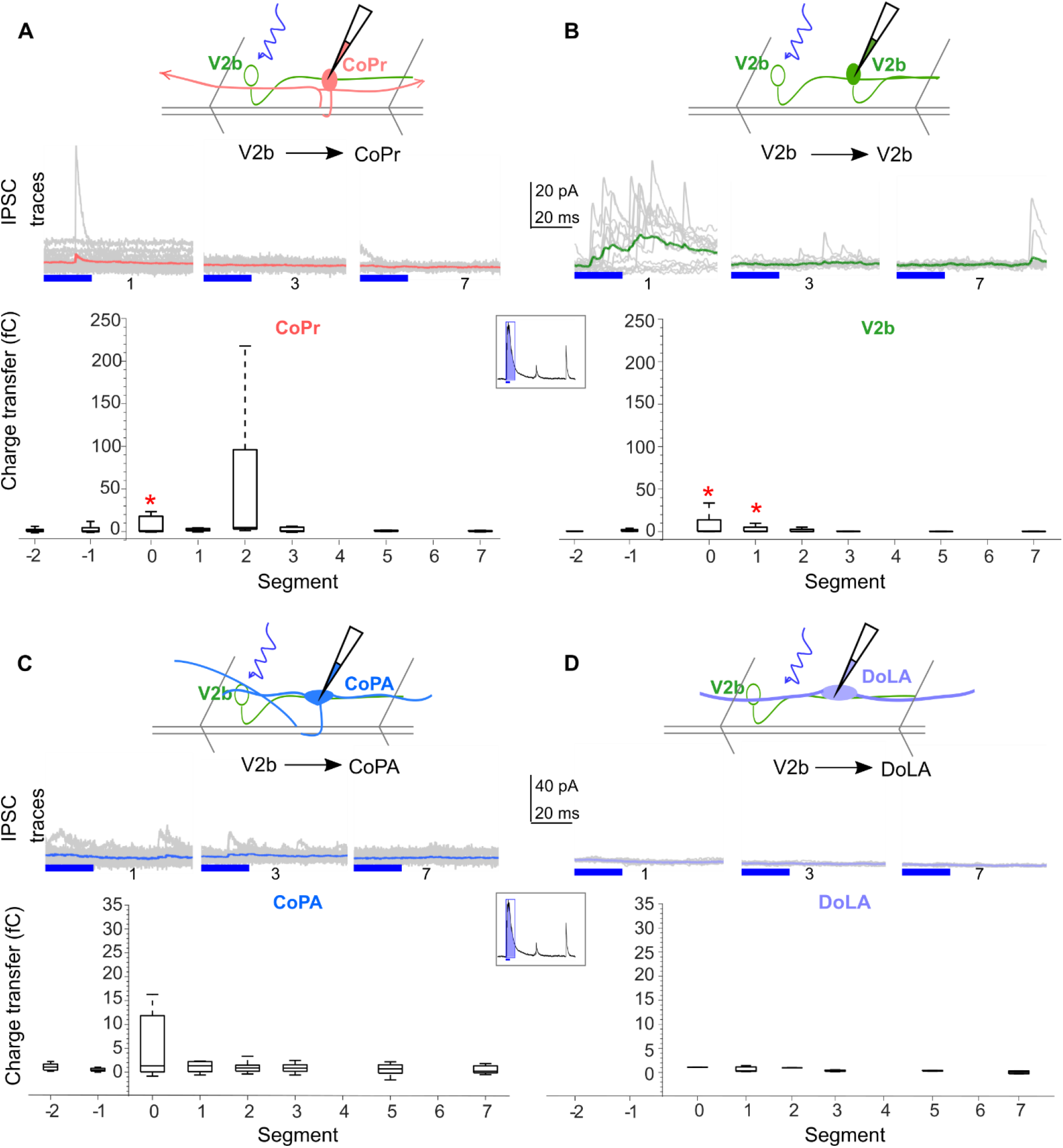
V2b inhibition to other spinal populations. A. V2b inhibition to commissural premotor neurons (CoPrs), an anatomically defined population likely including dI6 and V0 classes. Top: Schematic of the experimental design. Middle: Representative overlay of 15 traces recorded in a CoPr neuron during illumination of segments 1, 3, and 7 rostral to the recorded neuron soma. Colored trace represents mean. Duration of the optical stimulus is shown as a blue bar. Bottom: Box plot showing total charge transfer per segment (inset, illustration) recorded in V2a neurons. N = 5-7 neurons. B, C, D. Same as in A for V2b inhibition to other V2b neurons (B, N = 5-9 neurons), CoPA neurons (C, N = 11-12 neurons), and DoLA neurons (D, N = 3 neurons).

V2b neurons have been shown to inhibit each other^13^, but the rostrocaudal extent and amplitude of this inhibition is unknown. We observed V2b inhibition onto other V2b neurons exclusively locally (Fig. 6B, N= 5-9 neurons; Wilcoxon Sign Rank Test, *p* < 0.01), not at long range.

### V2b neurons do not inhibit two dorsal horn sensory populations

Finally, we wanted to test if V2b neurons target dorsal horn sensory neurons. In zebrafish, the Commissural Primary Ascending (CoPA) neurons, likely homologous to mammalian dI5 neurons, are glutamatergic neurons activated during the tactile reflex^53^. During swims, CoPAs are gated by V1 neurons^10,37^ and possibly others^53^. CoPA neurons are readily identifiable by their distinct axonal and dendritic morphology in posthoc dye fills. Fig. 6C shows representative traces (middle) and summary data (bottom) of V2b inputs to CoPA neurons. CoPAs did not receive any appreciable inhibition from V2b neurons (N= 11-12 neurons; Wilcoxon Sign Rank Test, *p* > 0.01) suggesting that V2b neurons are not involved in sensory gating of CoPA neurons during swims. Interestingly, CSF-cNs are known to contact CoPAs^30^. However, we did not capture this inhibition, in the caudal segments (Fig. 6C, Segments −1, −2), probably because CSF-cN inhibition to CoPA neurons is prominent slightly more distally, three segments away^30^. We also examined responses in Dorsal Longitudinal (DoLA) neurons, a sensory target likely homologous to mammalian dI4 neurons^54,55^. DoLAs are GABAergic and exhibit a characteristic axonal morphology but much less is known about their function. V2b neurons did not show any appreciable inputs to DoLA neurons (Fig. 6D; N= 3 neurons; Wilcoxon Sign Rank Test, *p* > 0.01). We conclude that inhibition of these sensory targets may not be a primary function of V2b neurons.

To compare V2b connectivity patterns across different cell types, we plotted a heat map showing median charge transfer for each population along the R-C axis (Fig. 7A). For each target neuron, charge transfer was first normalized to its total cellular conductance (inverse of input resistance) to allow a direct comparison across targets. This map clearly reveals in two key points: (i) Extensive V2b mediated inhibition of motor neurons and V2a neurons that slowly tapers over distance; and (ii) Selectively localized V2b-mediated inhibition onto V1, other v2b and commissural neurons. Taken together, this map shows that V2b inhibition is spatially structured in the rostrocaudal axis with distinct targets locally versus long-range.

**Figure 7:**
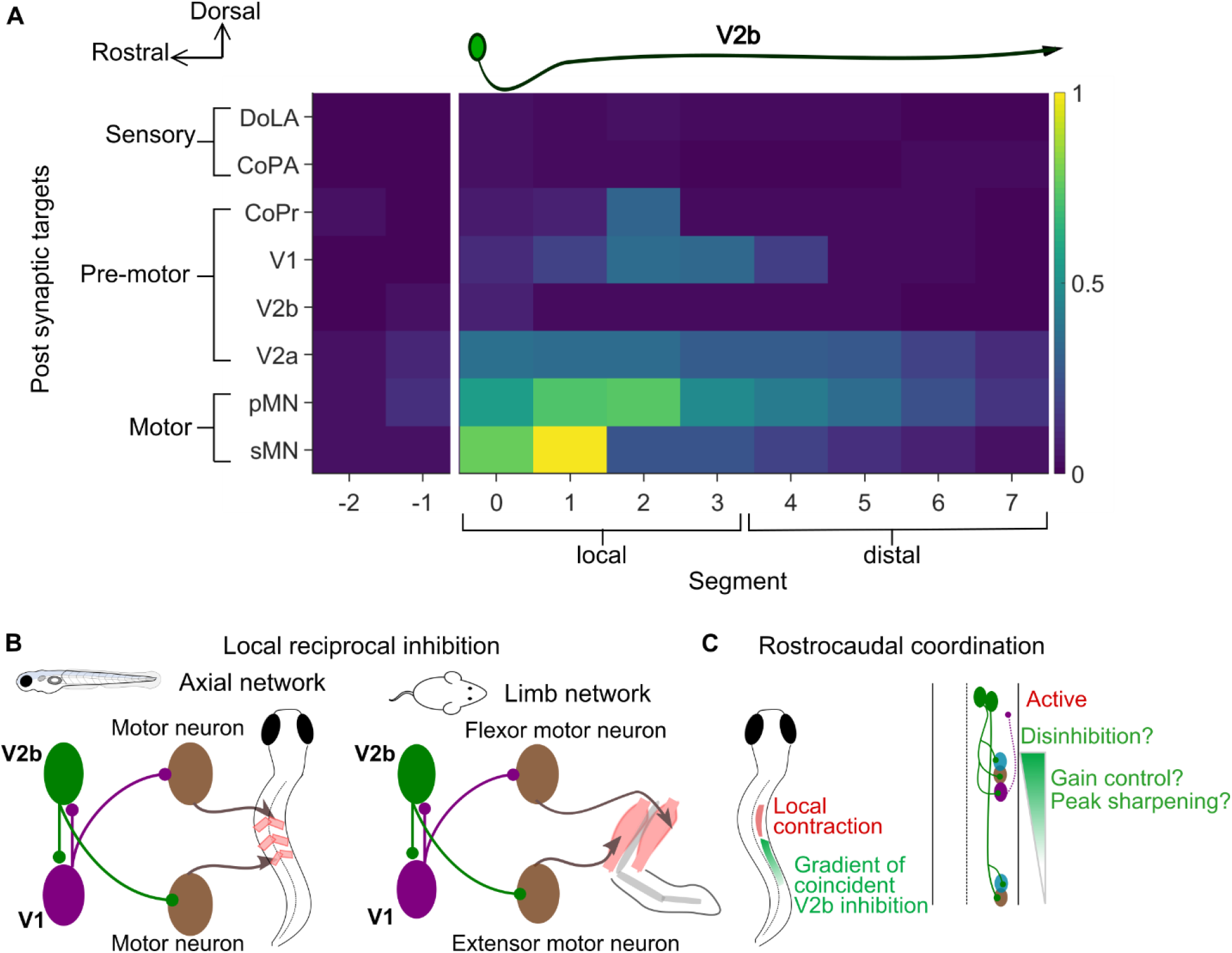
Summary of V2b connectivity. A. Heat map showing normalized charge transfer for the different post-synaptic targets along the rostro-caudal axis. The charge transfer per segment for each recorded neuronal target was normalized to its measured intrinsic neuronal conductance (inverse of R_in_). Median values of normalized charge transfer for each target cell population are plotted. Values for Segment 4 and Segment 6 were interpolated as averages of the two neighboring segments. The resulting values are plotted on the same color scale for all target populations. B. Schematic showing similarity of observed reciprocal inhibition between V1 and V2b neurons in axial circuits (left; this study and ref 10) and predicted reciprocal inhibition between V1 and V2b neurons in limb circuits (right). In both cases, there is a rostrocaudal asymmetry in connectivity. C. A proposed function of V2b neurons in rostrocaudal coordination. A gradient of V2b inhibition, arriving in phase with excitation to motor and V2a neurons might function to modulate gain, sharpen contraction and/or provide local disinhibition, facilitating rostrocaudal propagation.

## Discussion

In this study we show for the first time that V2b neurons are active largely in phase and leading the swim cycle, with heterogeneity across the population. V2b neurons inhibit motor neurons and several cardinal ventral horn spinal populations, including V1 and V2a neurons. In contrast, they do not inhibit two populations of dorsal horn sensory neurons, suggesting they have a predominantly motor role. Furthermore, V2b inhibition shows a selective spatial organization in the rostrocaudal axis, supplying long range inhibition to motor neurons and V2as but local connectivity to inhibitory V1 neurons, setting up a local disinhibitory circuit. In conjunction with previous work, this pattern of synaptic output indicates that axial V1 and V2b populations reciprocally inhibit each other with a rostrocaudal structure, similar to the inferred circuit for limb control in later vertebrates.

### Local reciprocal inhibition of V1 neurons

We previously demonstrated that V1 neurons inhibit the V2b population up to three segments rostrally^10^. Together with the V2b connectivity presented here, we conclude that V1 and V2b neurons reciprocally inhibit each other. Because our mapping was done at a population level, it is still unknown if single V1/V2b pairs show reciprocal inhibition. Nonetheless, this circuit motif closely resembles reciprocal inhibition for flexor-extensor modules (Fig. 7B). Mutual inhibition between Ia inhibitory interneurons has been reported in several species^24,40–42^ and is thought to be a crucial component for enforcing flexor-extensor alternation^56^. Though Ia inhibitory interneurons are composed of V1 and V2b populations, V1/V2b reciprocal innervation has so far not been systematically analyzed. We find that V1 and V2b neurons inhibit each other locally, up to three segments away. Specifically, V2b neurons inhibit V1 neurons located 2-3 segments caudally, whereas V1 neurons inhibit V2b neurons located 2-3 segments rostrally. This is consistent with the rostrocaudal and local structure of hindlimb flexor-extensor circuits^57^, where the predominant flexor motor neuron signal is at L2 (Lumbar 2), three segments rostral to the predominant extensor motor neuron signal at L5^16^. The origin of limb networks is debated. Some evidence suggests that tetrapod locomotion evolved from undulatory swimming and hence networks for limb control originated through gradual modification of circuits regulating axial musculature^5,58–61^. Our results showing V1/V2b reciprocally inhibitory circuits exist in axial networks may therefore provide evidence of an ancestral circuit motif that could have been co-opted for flexor-extensor control (Fig. 7B, left).

The functional relevance of V1/V2b reciprocal connectivity in axial motor control is still not clear. Interestingly, loss of V1 or V2b neurons in larval zebrafish shows opposing behavioral effects: loss of V1 neurons slows down locomotion^20^ while suppression of V2b neurons yields an increase in locomotor speed^13^. Current analysis of V1/V2b connectivity does not reveal any biases to any speed modules^10^. However, a closer look comparing inputs from V1 and V2b neurons to the same target populations reveal intriguing biases. For both primary motor neurons and V2a population, V2b inhibition is stronger than V1 inputs consistent with V2b function of slowing down locomotion. For secondary motor neurons, V1 inhibition is more profound than V2b, in agreement with V1 neurons facilitating fast locomotor speeds. This V1/V2b reciprocal inhibition and biased connectivity suggests this motif could have been utilized for implementing different speeds of locomotion in axial networks but remains to be explicitly tested.

### Coincident inhibition and rostrocaudal coordination

Locomotion requires efficient coordination along the rostro-caudal axis. In aquatic vertebrates, this takes the form of an S-shaped body during swimming, with a complete activity cycle from rostral to caudal^6^. Although waves of excitation drive motor neuron firing, precise control of timing may require coincident inhibition. Prior work concluded that V1 neurons provide a coincident inhibitory signal that helps terminate the locomotor burst, preventing extended firing^10,20^. One possible role for V2b neurons is an equivalent inhibitory signal on the rising phase of excitation (Fig. 7C). Our results showing that V2b neurons are active in phase or leading the swim cycle (Fig. 1) and that they provide a gradient of inhibition over long distances provides a potential mechanism for enforcing well-timed motor contractions in the ipsilateral axis.

Coincident inhibition is widespread in the cerebral cortex where it can impact multiple network parameters like synaptic gain, dynamic range, sharpening of sensory tuning and spike synchrony^62^. Motor neurons have been shown to receive coincident inhibition^63,64^ that modulates synaptic gain^65,66^, but the source of this inhibition is unknown. Interestingly, V1 neurons also provide on cycle inhibition^20^, but the timing of V1 spiking is substantially later than V2b activity, with peaks aligned with or after the peak of the motor burst.^20^ This phase relation allows V1 neurons to terminate motor bursts, permitting fast speeds of locomotion^20,38^. V2b neuron spiking, on the other hand, precedes the motor burst, coinciding with the rising phase of excitation. This temporal profile ideally positions V2b neurons for adjusting the gain of target neurons. The rostrocaudal structure of inhibition, tapering in strength at distal segments, is well suited to help sharpen the local wave of contraction (Fig. 7C).

It remains unclear whether the V2b neurons that did not fire at all during locomotion instead exhibit a specialized functional role, such as during postural or high-speed movements. It will be of interest to examine the firing properties of the different subtypes of V2b neurons^13^ across a wide range of motor outputs to determine their overall recruitment. The recorded V2b neuron activity on the rising phase of motor activity appears similar to the recruitment pattern of bifurcating (Type II) V2a neurons^31^, suggesting that these V2a neurons might provide a source of excitation to V2b neurons.

V2b neurons might also aid rostrocaudal coordination through an additional mechanism. Locomotion requires diagonal coupling of the left and right sides. In limbed vertebrates, the left forelimb and right hind limb are synchronously active. Modeling studies show that long range inhibition in the ipsilateral axis is crucial for this coupling^67^. Our results showing that V2b neurons selectively inhibit motor neurons and V2a neurons at long distances provide a potential template for enforcing long-range ipsilateral inhibition to facilitate diagonal coupling. This role of V2b neurons is indirectly supported in mice, where V2b neurons have been identified as a component of the long descending propriospinal neurons^14^ that facilitate interlimb coordination^15,68^. Extended V2b axonal projections covering half the spinal length, seen in mouse limb^14^ and midbody larval zebrafish^13^, could be a source for this long-range coordination.

Current models for generating rhythmic locomotion are often incomplete due to lack of knowledge on activity and connectivity of interneurons. While connections among and between several functional interneuron groups have been predicted in these models, a dearth of direct evidence makes linking these functions to known genetic classes difficult. In this study, by providing a detailed analysis of V2b activity during fictive locomotion and connectivity to motor and other spinal populations, we provide much needed information to fill gaps in current models and enhance our understanding of spinal motor circuits.

## Acknowledgments

This research was funded by the Pew Scholar Award (M.W.B), R01 DC016413 (M.W.B), a McKnight Scholar Award (M.W.B.), and the McDonnell Center for Cellular and Molecular Neurobiology Postdoctoral Fellowship 2021 (M.S.).

---

**Supplemental figure 1:**
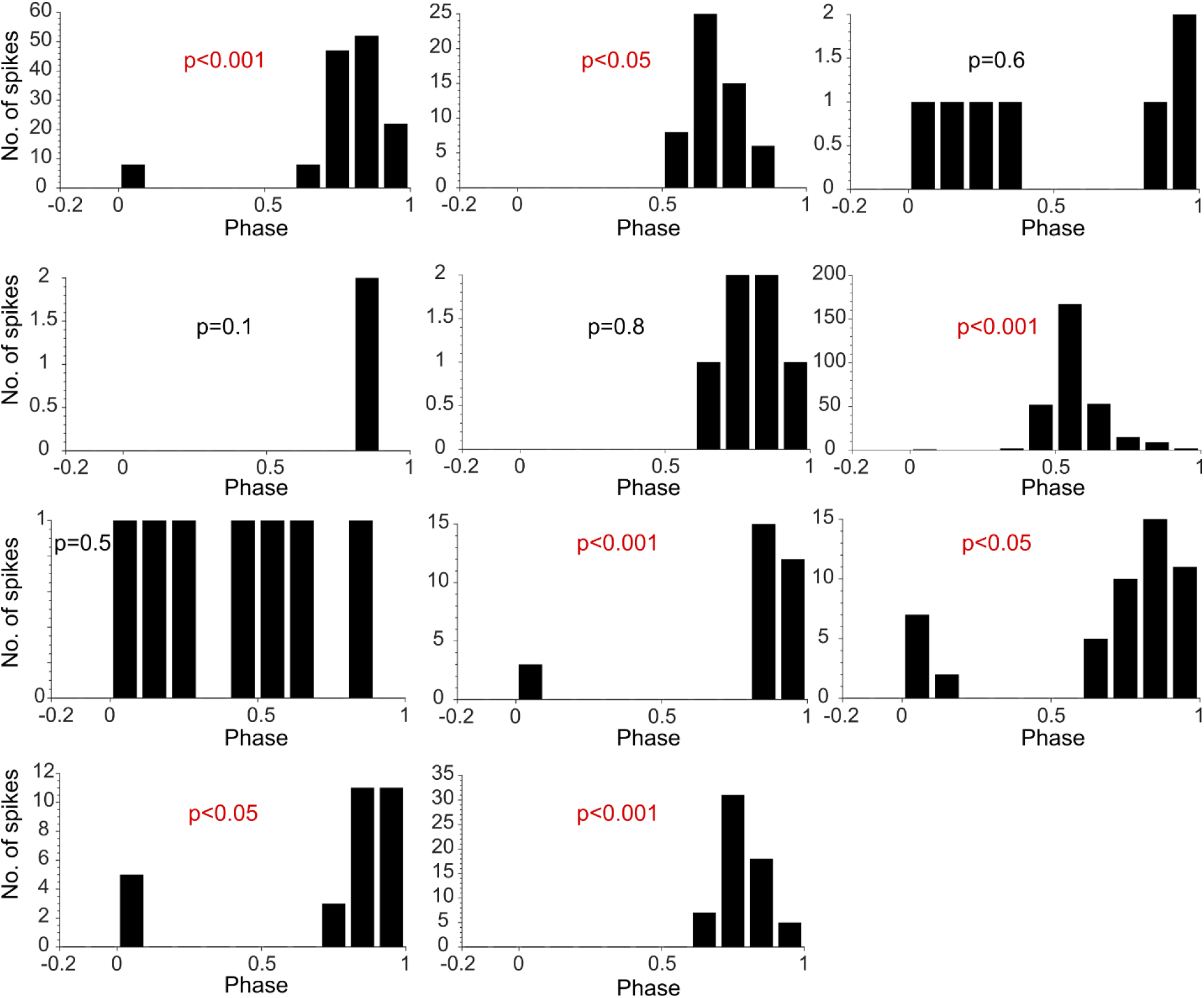
Phase relation of V2b neuron spiking to the swim cycle. Histogram showing spike timing of 11 different V2b neurons. Spike timing was normalized to phase of motor activity. Statistical significance was tested using Rayleigh’s Test for circular uniformity.

